# Gene expression of endangered coral (*Orbicella* spp.) in the Flower Garden Banks National Marine Sanctuary after Hurricane Harvey

**DOI:** 10.1101/703447

**Authors:** Rachel M. Wright, Adrienne M.S. Correa, Lucinda A. Quigley, Sarah W. Davies

## Abstract

About 160 km south of the Texas–Louisiana border, the East and West Flower Garden Banks (FGB) have maintained >50% coral cover with infrequent and minor incidents of disease or bleaching since monitoring began in the 1970s. However, a storm that generated coastal flooding, which ultimately interacted with the reef system, triggered a mortality event in 2016 that killed 2.6% of the East FGB. To capture the immediate effects of storm-driven freshwater runoff on coral and symbiont physiology, we leveraged the heavy rainfall associated with Hurricane Harvey in late August 2017 by sampling FGB corals at two times: September 2017, when salinity was reduced; and one month later when salinity had returned to typical levels (~36 ppt in October 2017). Tissue samples (N = 47) collected midday were immediately preserved for gene expression profiling from two congeneric coral species (*Orbicella faveolata* and *Orbicella franksi*) from the East and West FGB to determine the physiological consequences of storm-derived runoff. In the coral, differences between host species and sampling time points accounted for the majority of differentially expressed genes. Gene ontology enrichment for genes differentially expressed immediately after Hurricane Harvey indicated increases in cellular oxidative stress responses. Although tissue loss was not observed on FGB reefs following Hurricane Harvey, our results suggest that poor water quality following this storm caused FGB corals to experience sub-lethal stress. We also found dramatic expression differences across sampling time points in the coral’s algal symbiont, *Breviolum minutum*. Some of these differentially expressed genes may be involved in the symbionts’ response to changing environments, whereas a group of differentially expressed post-transcriptional RNA modification genes also suggest a critical role of post-transcriptional processing in symbiont acclimatization. In this study, we cannot disentangle the effects of reduced salinity from the collection time point, so these expression patterns may also be related to seasonality. These findings highlight the urgent need for continued monitoring of these reef systems to establish a baseline for gene expression of healthy corals in the FGB system across seasons, as well as the need for integrated solutions to manage stormwater runoff in the Gulf of Mexico.

## Introduction

Hurricane Harvey intensified to a Category 4 storm over the Gulf of Mexico on 24 August 2017 and made landfall on the Texas coast shortly after. The storm stalled over land for several days, resulting in an estimated 33 trillion gallons of rainfall and more than 100,000 damaged homes in Texas and Louisiana (van Oldenborgh et al. 2017; Shultz and Galea 2017). In the weeks following the storm’s retreat, Galveston Bay experienced heavy freshwater outflow, which elevated sea levels for more than four days and reduced salinity to nearly zero at multiple monitoring stations in the bay (Du et al. 2019). Besides being hyposaline, this storm-derived water mass contained high nutrients levels and other compounds of terrigenous origin, which had the potential to shift pelagic communities and their associated processes (Lefebure et al. 2013; Jonsson et al. 2017), and to impact the health of organisms that came in contact with this mass (Liñán-Cabello et al. 2016).

The Flower Garden Banks National Marine Sanctuary (FGBNMS), which harbors one of the few remaining reef systems in the wider Caribbean with >50% coral cover (Johnston et al. 2016; Gardner et al. 2003), sits southeast of Galveston Bay and thus risked exposure to the offshore water mass generated by Hurricane Harvey. Impacts from this water mass were of particular concern given that approximately one year earlier (July 2016) a portion of the East Flower Garden Bank (EFGB) experienced a highly localized die-off event over 5.6 ha of reef (2.6% of the site), which contributed to partial or full mortality of an estimated 82% of coral colonies and declines in many other benthic invertebrates (Johnston et al. 2019). While no abiotic data were collected at the EFGB mortality site during the 2016 die-off, salinity and temperature measurements from nearby regions around the time of the event suggest low dissolved oxygen played a critical role in this mortality event (Kealoha et al. *in revision*). Heavy rainfall along the coast immediately before the 2016 mortality event resulted in unusually high levels of freshwater runoff, which likely contributed to water stratification and reduced gas exchange at the affected site (Kealoha et al. *in revision*). Reef-building corals are among the tropical marine species most vulnerable to the effects of hurricanes (Woodley et al. 1981; Gardner et al. 2005). Coral colonies can be impacted by hurricanes via physical damage from waves (Highsmith, Riggs, and D’Antonio 1980; Bries, Debrot, and Meyer 2004), smothering by sediments (Highsmith, Riggs, and D’Antonio 1980; Bries, Debrot, and Meyer 2004), and reductions in water quality (*e.g*., Manzello et al 2013; Edmunds et al. 2019, Nelson & Altieri 2019). Poor water quality can harm corals through a variety of mechanisms including reduced salinity and aragonite saturation state and increased turbidity, nutrients, and pollutants. Low salinity caused by heavy rainfall associated with extreme storms can trigger mass loss of the algal endosymbionts of corals (Family Symbiodiniaceae; Goreau 1964; Bries, Debrot, and Meyer 2004; Pengsakun et al. 2019). For example, rainfall from Tropical Storm Isaac in 2012 caused a week-long reduction in aragonite saturation along the Florida Keys reef tract, which resulted in reduced coral calcification (Manzello et al. 2013). Turbidity increases due to terrestrial runoff during storms can also significantly reduce light penetration over reefs reducing the ability of the algal symbiont to photosynthesize. For example, Hurricanes Irma and Maria caused temporary periods of complete darkness at a depth of 19 m on a reef off St. John in the US Virgin Islands in 2017 (Edmunds et al. 2019). As a result, these storms caused a 20% reduction in daily integrated underwater photosynthetic photon flux density over a period of 69 days. Terrestrial runoff following extreme storms can also increase nutrient levels in these normally oligotrophic reef-associated waters, causing bacterial blooms that can ultimately trigger oxygen drawdown and suffocation of reef organisms (*e.g*., Kealoha *et al*. *in revision*, Nelson and Altieri 2019). An overview of the diverse mechanisms by which shifts in water quality attributes can trigger low dissolved oxygen conditions is provided in Nelson & Altieri (2019).

Given the FGB’s recent history of coral mortality following an extreme storm event, the aim of this study was to identify the immediate physiological impacts of a major storm (Hurricane Harvey, 2017) in the same reef system: the East and West FGB. Two congeneric coral species (*Orbicella faveolata* and *Orbicella franksi*) were sampled at two time points: immediately after Hurricane Harvey in September 2017 and one month later. Global gene expression profiling of these corals and their photosynthetic algal symbionts (*Breviolum minutum*) was conducted to determine the physiological consequences of a water mass generated by an extreme storm on dominant reef-building coral species in the FGB.

## Materials and Methods

### Pelagic Water Properties after Hurricane Harvey

Water properties, including salinity (ppt) and temperature (°C), were measured by the Texas Automated Buoy System Real Time Ocean Observations, Buoy V (27° 53.7960’N, 93° 35.8380’W; sensor depth 2m), before and after coral sampling. Buoy V is approximately 3km from the EFGB and 25km from the WFGB). Data were downloaded from the archives: http://tabs.gerg.tamu.edu/tglo/tabsqueryform.php?buoy=V. Unfortunately, water property data at Tabs Buoy V do not exist for much of August 2017, including when Hurricane Harvey formed over the Gulf of Mexico (approximately 25 August 2017). In the days prior to the September coral collection, surface salinity and temperature were reduced (leftmost asterisks in Fig. 1), suggesting anomalous freshwater runoff effects from the storm. Surface salinity recovered by the second collection time point (October 2017; Fig. 1, top right asterisk). Henceforth, we refer to the first sampling time point (September 2017) as “sub-lethal stress” and the second sampling time point (October 2017) as “recovery” to describe the hypothesized effects of the storm on the coral and its algal symbiont at those times.

**Figure 1:**
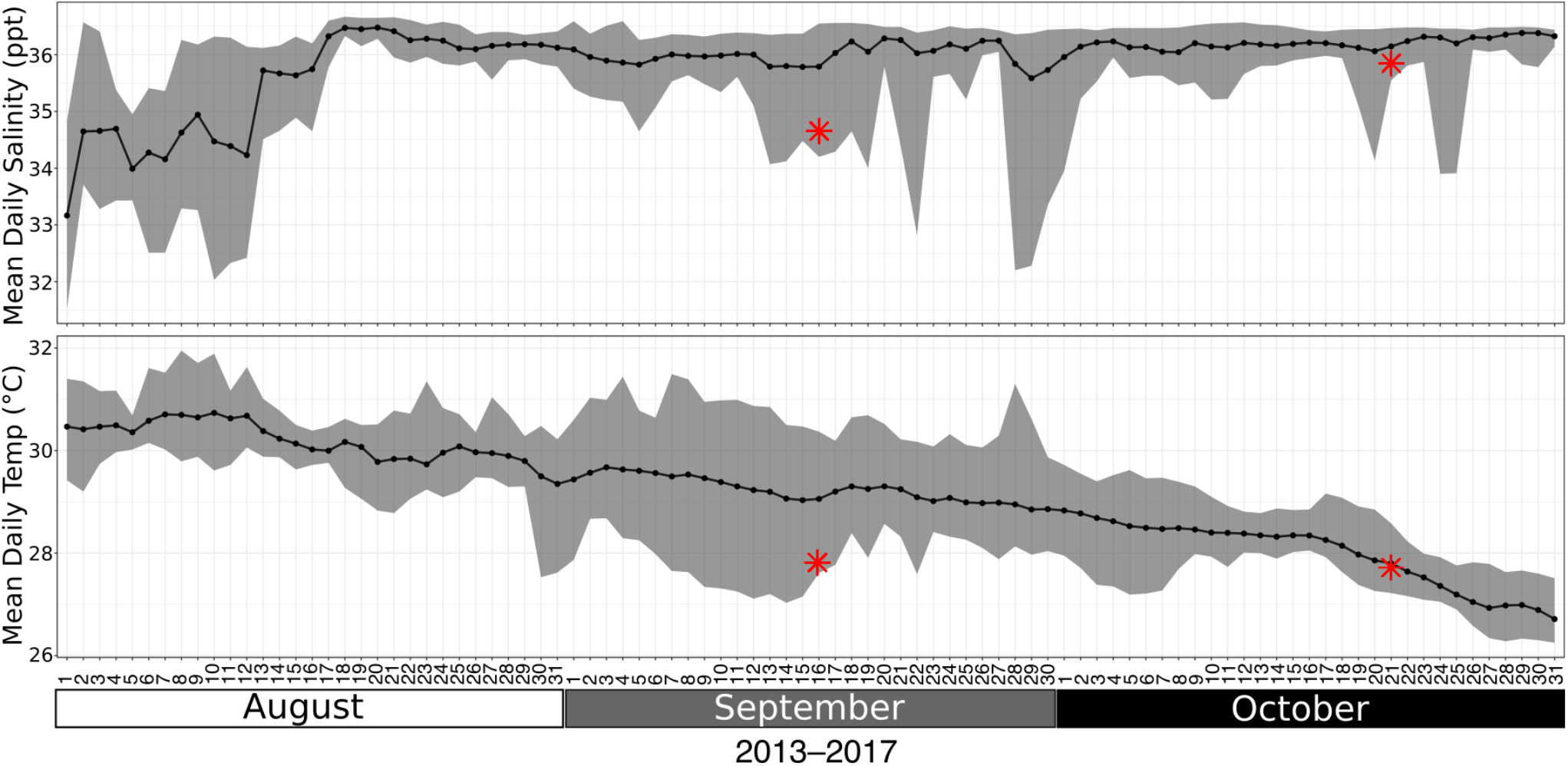
Salinity (ppt, top) and temperature (°C, bottom) at buoy V near the coral sampling sites at the East and West Flower Garden Banks for the weeks surrounding the time of sampling. Black circles represent daily means from 2013–2017. Grey ribbons encompass minimum and maximum values from 2013–2017. Red asterisks represent mean daily values on September 16, 2017, the first collection time point, and on October 21, 2017, the first day of the second coral collection.

### Coral Collections

Tissue fragments were collected from individually tagged *Orbicella faveolata* and *Orbicella franksi* coral colonies in the FGBNMS (northwest Gulf of Mexico) during periods of “sub-lethal stress” (on 16 September 2017, EFGB only) and “recovery” (21-24 October 2017, East and West Banks, Supp. Mat. Table 1). In total, 23 samples of *O. faveolata* and 24 samples of *O. franksi* were collected over the two sampling periods (Table 1).

**Table 1:**
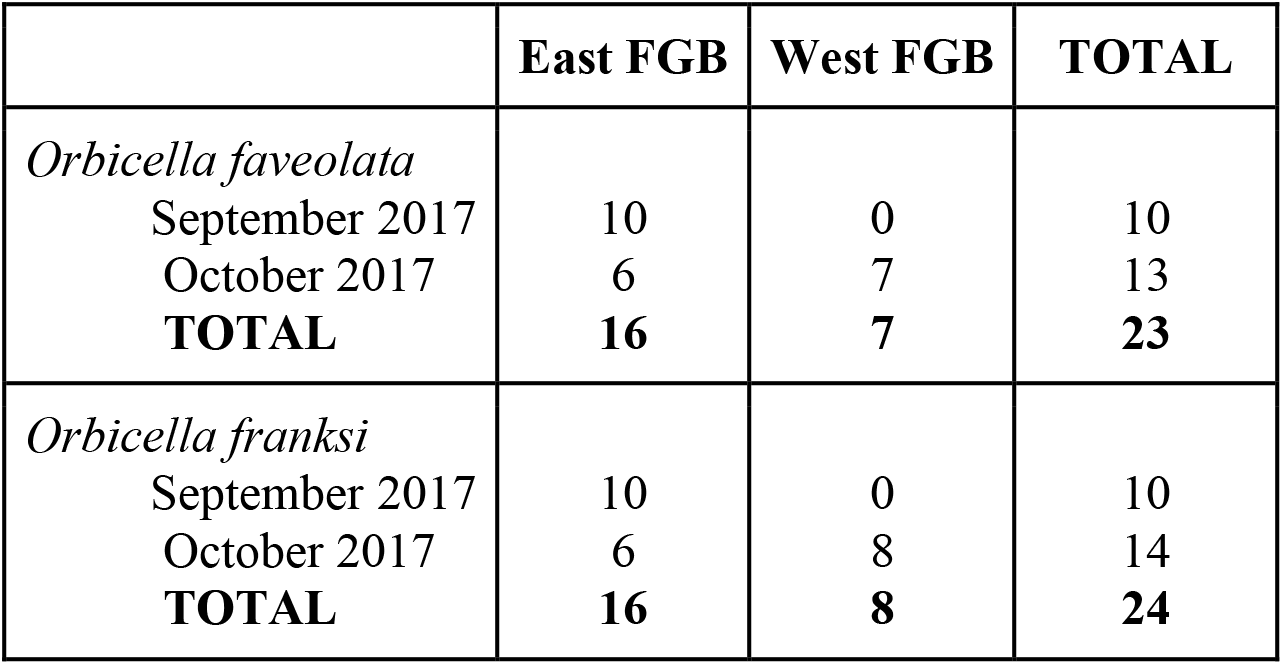
Overview by species, location and time of the 47 coral colonies sampled from the East and West Banks of the Flower Garden Banks National Marine Sanctuary (northwest Gulf of Mexico) in this study.

The depths of sampled colonies ranged from 19.2 to 24.1m. Details on the locations and depths of samples collected for each coral species are provided in Supp. Mat. Table 1. Samples were collected from the tops of colonies using a hammer and a species-specific chisel and were immediately placed in pre-labeled upside-down 15mL falcon tubes containing 200 proof molecular grade EtOH free of air bubbles.

### Gene Expression Library Preparation

RNA was isolated from 47 coral tissue samples using the RNAqueous-Micro Total RNA Isolation Kit (Invitrogen). Coral fragments were placed in tubes containing 150 μL of lysis buffer and glass beads (Sigma, 150–212μm). Samples were placed in a bead blaster at 5 m/s for 1 minute and then centrifuged at a speed of 16.5 ×g for 1 minute. The supernatant was transferred to a new tube and centrifuged again at 16.5 ×g for 2 minutes, and then transferred to a final tube. RNA was eluted, washed, and DNased using 10× DNase I. First-strand synthesis, cDNA amplification, barcoding, and pooling were performed according to an established protocol (Meyer, Aglyamova, and Matz, 2011; Dixon et al 2015). A total of 47 gene expression libraries were prepared, with 20 from September 2017 and 27 from October 2017. Sequenced reads have been uploaded to the National Center for Biotechnology Information Short Read Archive under accession number PRJNA552981.

### Gene Expression Analysis

Adapter sequences were trimmed and low quality reads (minimum quality score = 20; minimum percent bases above minimum quality score = 90%) were filtered using FASTX tools (Hannon 2010). Reads were mapped to a composite coral host and algal symbiont transcriptome, which included concatenated sequences from the coral, *Orbicella faveolata* (Pinzon et al. 2015), and its algal symbiont, *Breviolum minutum* (Parkinson et al. 2016), using Bowtie 2 (Langmead and Salzberg 2012). Given that only *B. minutum* and its haplotypes have been reported from FGBNMS *O. faveolata* and *O. franksi* to date (Santos et al. 2006, Green et al. 2014, Correa et al. *in prep.*), this species was the only algal reference transcriptome used. Statistical analyses were conducted in R version 3.4.0 (R Core Team 2017). Isogroups (henceforth called “genes”) with a base mean < 3 across all samples were removed from the analysis. Expression sample outliers were detected using *arrayQualityMetrics* (Kauffmann, Gentleman, and Huber 2009). Differentially expressed genes (DEGs) were identified using *DESeq2* (Love, Huber, and Anders 2014). Wald tests were performed to calculate contrasts between sampling time points, host species, and collection location (i.e., EFGB or WFGB). Log-fold change (LFC) values for sampling time point are expressed relative to the October collection (*e.g*., negative LFC indicates upregulation in September relative to October or, equivalently, downregulation in October relative to September). False-discovery rate (FDR) p-values were adjusted using the Benjamini–Hochberg procedure (Benjamini and Hochberg 1995). To test for overall transcriptomic differences across samples, principal coordinate analysis and permutational analysis of variance testing on Manhattan dissimilarity matrices was performed using *vegan* (Dixon 2003). Gene expression heat maps were generated using *pheatmaps* (Kolde 2012) and gene ontology enrichment was performed based on signed adjusted p-values using *GO-MWU* (Wright et al. 2015).

## Results

### Gene Expression Associated with Coral Host, Sampling Time Point, and Collection Site

An average of 1.85 × 10^6^ host reads per sample and 0.2 × 10^6^ symbiont reads per sample remained after quality filtering and mapping to the *O. faveolata* and *B. minutum* transcriptomes. Both coral host species were mapped to the *O. faveolata* transcriptome, but we observed no significant difference in mapping efficiency between *O. faveolata* (50.7±7.9%) and *O. franksi* (47.0±10.3%, p = 0.179). In the coral host, differences between *Orbicella* species (p = 0.008) and sampling time points (p = 0.026) explain the majority of the observed differences in gene expression profiles (Fig. 2A). There was no significant variance in host gene expression associated with sampling locations (p = 0.063). *Breviolum minutum* expression profiles were impacted by sampling time point (p = 0.007), but not host species (p = 0.366) or sampling location (p = 0.125) (Fig. 2B).

We used an adjusted p-value threshold of 0.05 calculated by the Wald test to identify significantly Differentially Expressed Genes (DEGs) in the coral host (Fig. 3A) and algal symbiont, *Breviolum minutum* (Fig. 3B). In the coral host, we identified 769 (3.9% of transcriptome) and 265 (1.3% of transcriptome) DEGs when comparing between species (*O. faveolata vs*. *O. franksi*) and time (September sub-lethal stress *vs.* October recovery), respectively. Fifteen genes were significantly differentially expressed between the EFGB and WFGB sampling locations in the coral host (0.08% of transcriptome). In *B. minutum*, we identified 1,471 (4.6% of transcriptome) and 21 (0.07% of transcriptome) DEGs when comparing between time (September sub-lethal stress *vs.* October recovery) and sampling location (EFGB *vs*. WFGB), respectively. We found only two DEGs when comparing *B. minutum* expression between the two coral host species (<0.01% of transcriptome).

**Figure 2:**
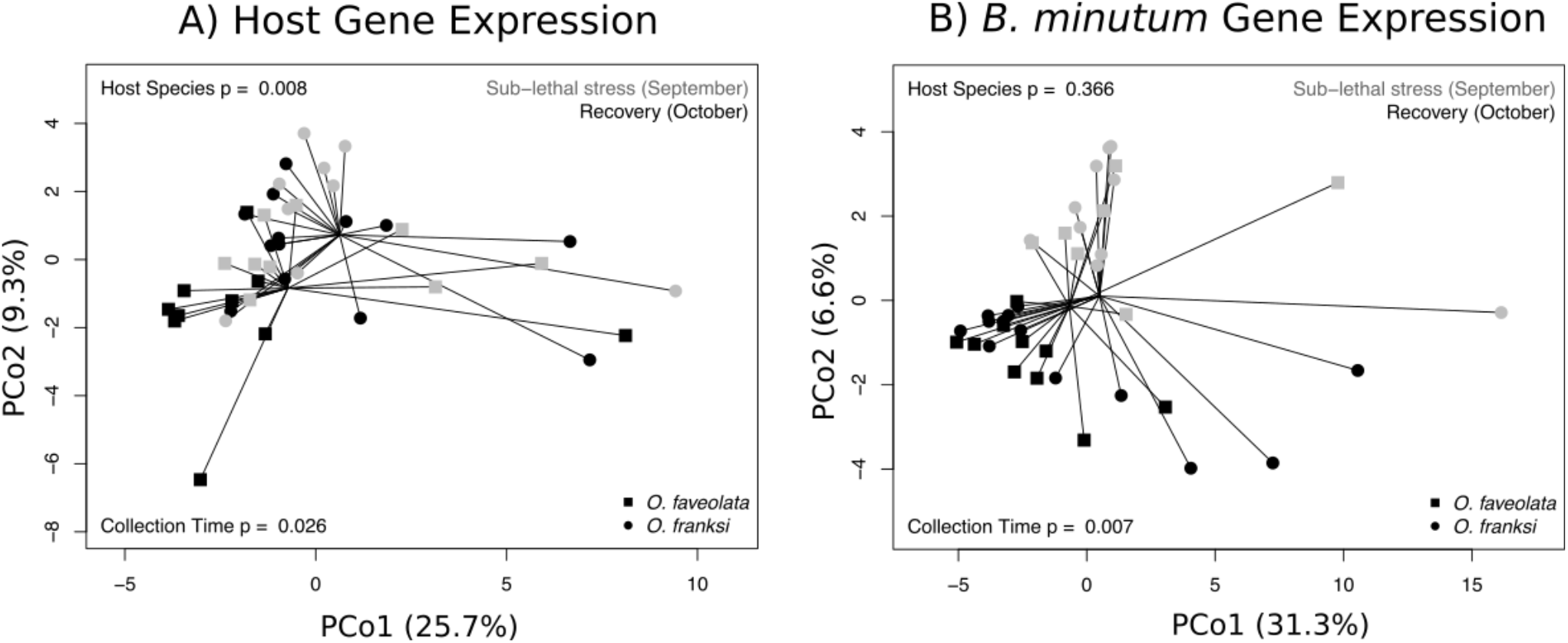
Principal coordinates analysis for coral host (**A**) and *B. minutum* (**B**) gene expression profiles. Each point is a sample and points closer together exhibit more similar expression profiles. Spider lines connect samples originating from the same coral species (*Orbicella faveolata* or *Orbicella franksi*), as indicated. Colors indicate whether the sample was collected during the sub-lethal stress (September, grey) or recovery (October, black) period. P-values were generated by permutational multivariate analysis of variance using distance matrices.

**Figure 3:**
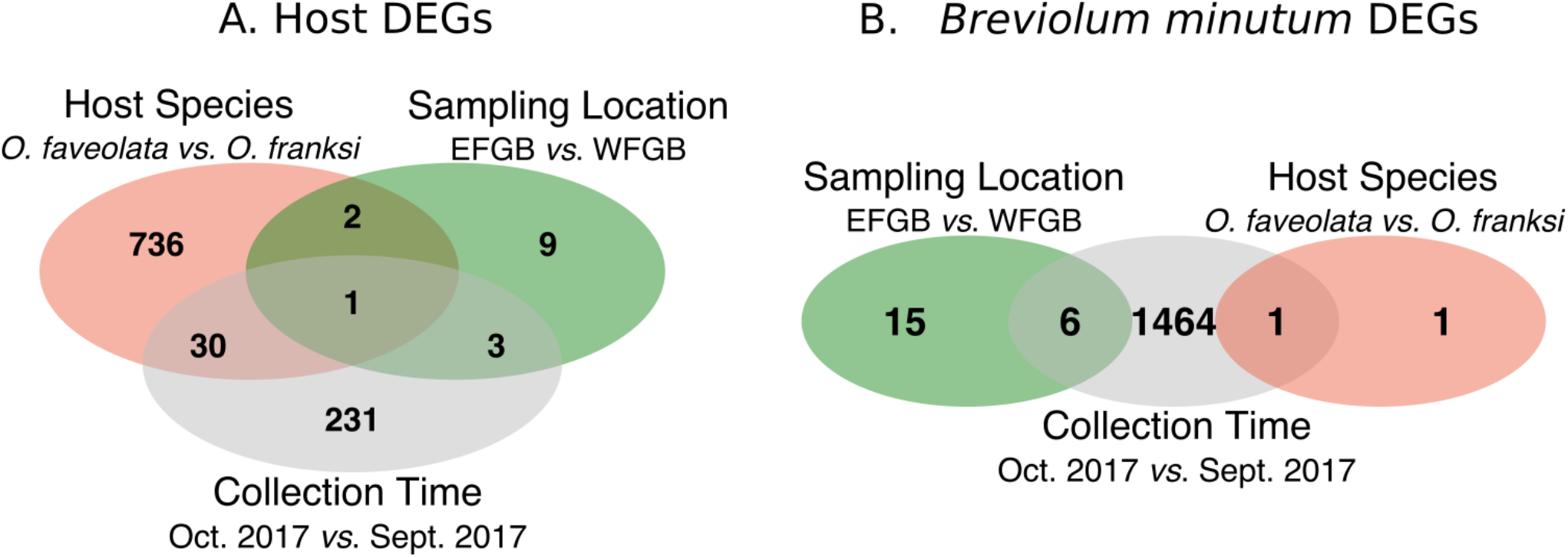
Venn-diagrams specifying the number of unique and shared differentially expressed genes (adjusted p-value < 0.05) by *Orbicella* host species, sampling location, and collection time point (September sub-lethal stress and October recovery) in the coral hosts (**A**) and *Breviolum minutum* (**B**).

### Gene Ontology Enrichment

In the coral host, gene ontology (GO) categories enriched during the sub-lethal low salinity stress event (September 2017) included antioxidant activity (p = 0.013), cell redox homeostasis (p = 0.039), and mitochondrial membrane parts (p = 4.04e-7) (Supp. Mat. Table 2). When normal salinity levels had returned, during the ‘recovery’ time point, many categories related to growth and cellular propagation were enriched within up-regulated genes, such as cell division (p = 0.003) and organelle fission (p=0.003). No GO terms were significantly enriched when comparing genes differentially expressed by either host species or sampling location.

The annotated coral host genes differentially regulated across the sub-lethal stress and recovery periods are shown in Figure 4. Relative to the recovery period, corals under sub-lethal low salinity stress up-regulated small cysteine rich protein 4 (LFC = −3.48, FDR = 4.2e-8) and down-regulated protein WNT-5 (LFC = 1.76, FDR = 0.006).

**Figure 4:**
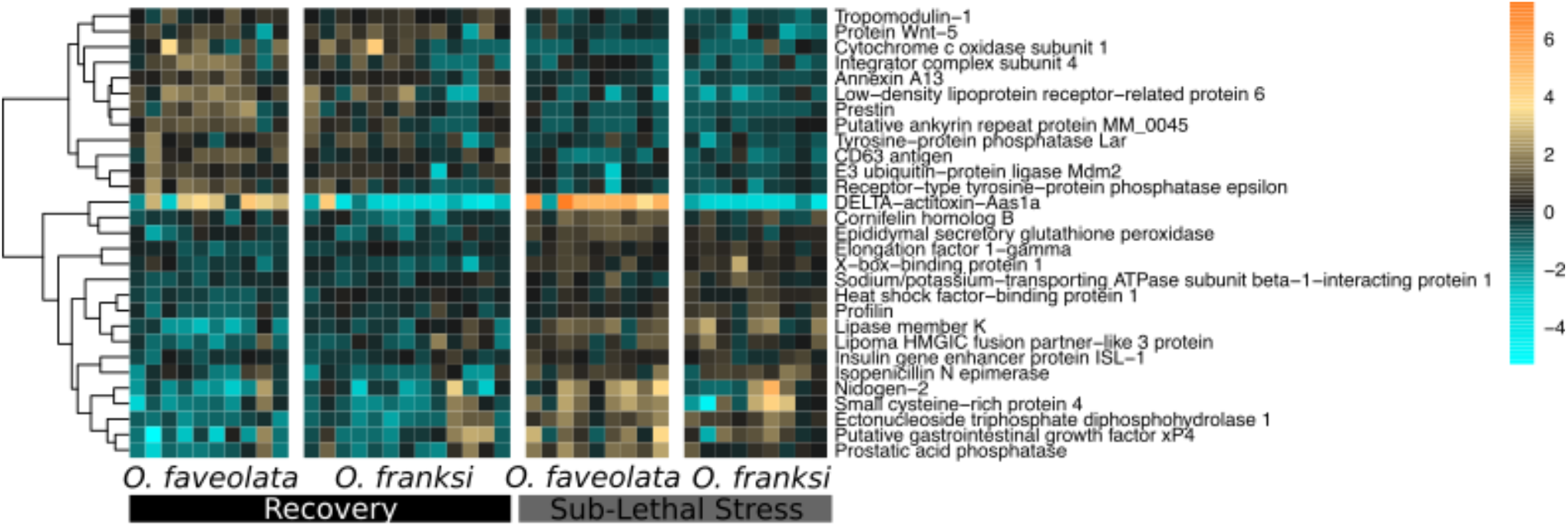
*Orbicella* spp. host gene expression differences across collection time points: sub-lethal stress *vs.* recovery (September 2017 *vs.* October 2017, FDR < 0.01). Rows are genes and columns are samples. The color scale indicates log2-fold change relative to the mean expression of each gene across all samples. Genes are hierarchically clustered based on Pearson’s correlations of expression across samples. Coral species are indicated below each column. Rectangles below coral species labels indicate the collection time point (black = October 2017 recovery; grey = September 2017 sub-lethal stress).

In the algal symbiont, *B. minutum*, enriched GO categories under sub-lethal low salinity stress conditions (September 2017) included oxidoreductase activity (p = 0.03) and transmembrane transport (p = 2.76e-5) (Supp. Mat. Table 3). When average salinity levels had returned in October 2017 (i.e., during ‘recovery’), many GO categories related to DNA replication and RNA splicing were enriched with up-regulated genes (Supp. Mat. Table 3). RNA splicing was also enriched in algal symbionts hosted by *O. faveolata* relative to algal symbionts hosted by *O. franksi* (Supp. Mat. Table 3).

In *Breviolum minutum*, the majority of significant DEGs were upregulated during the recovery period (October 2017) or, equivalently, downregulated during the sub-lethal salinity stress event in September 2017 (Figure 5). These genes include an RNA helicase (LFC = 2.0, FDR = 4.85e-5) and a nonsense-mediated mRNA decay protein (LFC = 1.5, FDR = 1.80e-5).

**Figure 5:**
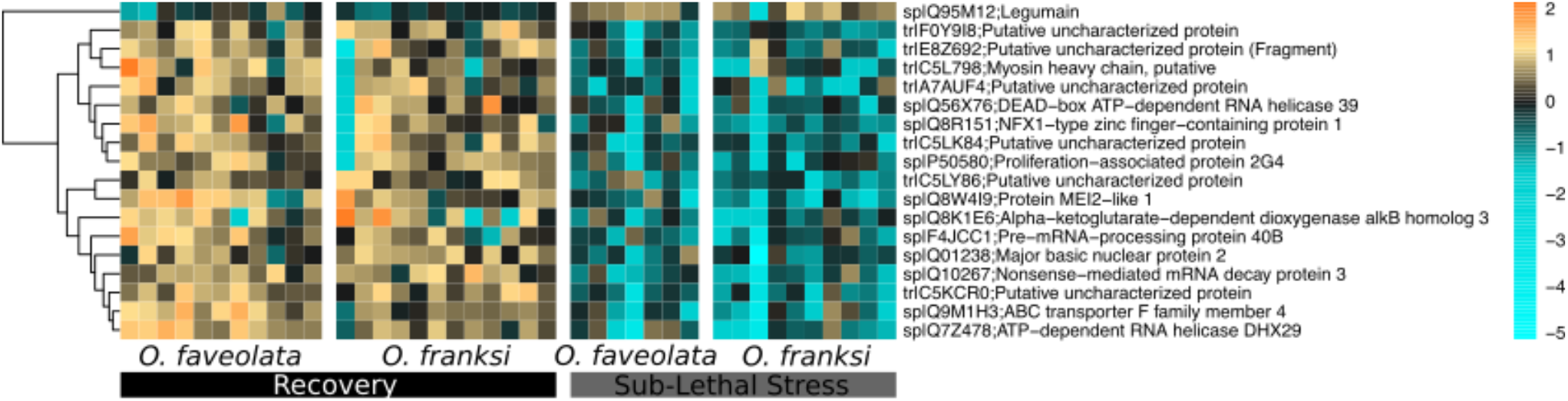
*Breviolum minutum* gene expression differences across sampling time: sub-lethal stress *vs.* recovery (September 2017 *vs.* October 2017, FDR < 0.0001). Rows are genes and columns are samples. The color scale indicates log2-fold change relative to the mean expression of each gene across all samples. Genes are hierarchically clustered based on Pearson’s correlations of expression across samples. Coral species are indicated below each column. Rectangles below coral species labels indicate the collection time point (black = October 2017 recovery; grey = September 2017 sub-lethal stress.

## Discussion

### Transcriptomic Effects of Freshwater from Hurricane Harvey on Coral Holobionts

The objective of this study was to use global gene expression profiling to determine the effects of a low salinity event associated with freshwater from Hurricane Harvey on two species of coral hosts and their algal symbionts in the Flower Garden Banks (FGB) National Marine Sanctuary (northwest Gulf of Mexico). Based on differences in pelagic water parameter data and coral holobiont gene expression differences between the two sampled time periods, we interpret that FGB coral and their symbionts were exhibiting early signs of low salinity stress as a result of freshwater runoff in September 2017, but that the influence of this freshwater influx was alleviated (*i.e*. recovered) by late October 2017.

#### Oxidative stress in the coral host

Corals are osmoconformers: when exposed to hyposaline conditions, water flows into their cells, thereby reducing internal cellular osmotic pressure (Titlyanov et al. 2000). The amount of damage cells sustain under reduced salinity depends on the extent of this osmotic pressure reduction and the length of time that cells are exposed to the stress (Berkelmans et al. 2012). *Stylophora pistillata* fragments exposed to five salinity concentrations ranging from 20–32 ppt showed increasingly severe cellular pathologies, including cell swelling and symbiont expulsion, with decreasing salinity (Downs et al. 2009). Within the coral cell, osmotic changes disrupt electron transport at mitochondrial membranes and increase reactive oxygen species produced by mitochondria. Consequently, robust mitochondrial antioxidant function is thought to be a major determinant of cellular management of osmotic stress (Pastor et al. 2009).

In both coral host species, GO categories were enriched with genes involved in antioxidant activity and mitochondrial structural components, suggesting the presence of oxidative damage that may compromise mitochondrial function (Supp. Mat. Table 2). A recent experimental study in *Acropora millepora*, a reef-building coral in the Great Barrier Reef, also found a strong antioxidant response to reduced salinity (Aguilar et al. 2019), suggesting that this response mechanism is conserved across coral genera. The upregulation of antioxidant-encoding genes has been described in corals exposed to a variety of biotic and abiotic threats, including increased temperature (Barshis et al. 2013; Dixon et al. 2015), acidification (Davies et al. 2016; Lopes et al. 2018), and disease (Wright et al. 2017, 2015; Daniels et al. 2015).

Given the diversity of stressors that trigger redox responses in corals, sub-lethal oxidative damage resulting from one stressor may contribute to coral declines when multiple threats occur simultaneously or successively (Carilli et al. 2009; Adjeroud et al. 2009). This study adds to the large body of evidence that genetic markers for antioxidant capability may be useful reef management tools to monitor coral health in the face of multiple climate change-related stressors (Jin et al. 2016).

#### *Expression responses in* B. minutum

At the FGB, both *Orbicella* species investigated here have been found to exclusively host *B. minutum* (Santos and LaJeunesse 2006; Green et al. 2014), Correa et al. *in prep.*), although subtle haplotype differences within *B. minutum* have been detected between these two host species as well as between the east and west FGB (Green et al. 2014). When comparing gene expression of *B. minutum*, we detected no major differences in expression between coral host species or sampling location and instead we found a significant effect of sampling time point on expression (Fig. 2B, Fig. 5). This gene expression response is in contrast to previous studies investigating the effects of multiple stressors on algal symbiont gene expression, which generally detect a paucity of expression changes and these changes are muted relative to their coral hosts (Barshis et al. 2014; Leggat et al. 2011; Davies et al. 2018, but see Baumgarten et al. 2013). One potential explanation is that we sampled before host buffering or acclimatization mechanisms diminished the symbiont response (*e.g*., Takahashi et al. 2013; Maboloc et al. 2015) or that our findings are primarily driven by seasonal fluctuations (Brown et al. 1999), which have not been explicitly characterized in Symbiodiniaceae *in hospite* to our knowledge.

A major category of genes differentially expressed in *B. minutum* across time points is associated with RNA-modification (Supp. Table 3; Fig 5). These candidates include a gene encoding a nonsense-mediated RNA decay protein (LFC = 1.5, FDR = 1.8e-5; Figure 5) and a gene encoding Regulator of Nonsense Transcripts 1 homolog (sp|Q9HEH1, LFC = 1.7, FDR = 0.03), which were both downregulated in September 2017 during the storm-induced low salinity period. In plants and mammals, nonsense-mediated decay (NMD) is inhibited during stress to allow proper activation of stress response functions. For example, inhibition of NMD under hypoxia augments the cellular stress response in mammalian cells (Gardner 2008) and inhibition of NMD in plant cells under pathogen attack stimulates plant defenses (reviewed in Shaul 2015). In our study, the downregulation of NMD-related genes may indicate symbiont stress sustained as a result of hyposalinity. Furthermore, a gene encoding Regulator of Nonsense Transcripts 1 homolog was also found to be differentially expressed in another coral symbiont, *Durusdinium* (formerly *Symbiodinium*) *trenchii*, within a juvenile *Acropora tenuis* host under benign conditions (Yuyama et al. 2018), suggesting that the gene product may play a role in normal interactions between the coral host and algal symbiont.

While these gene expression differences may be responses to the sub-lethal low salinity stress event associated with Hurricane Harvey experienced in September 2017, we also cannot disentangle responses to this event from seasonal changes occurring between the sampling time points, which would include lower light levels associated with slightly shorter and cooler days on average (Figure 1). Based on previous experimental studies conducted in other systems, the salinities observed at the FGBNMS in September 2017 may not have been low enough to trigger a response in the symbiont. Experimental exposures to low salinity (15–33.5 ppt) in *S. pistillata* caused symbiont loss coincident with reductions in photosynthetic efficiency (Kerswell and Jones 2003). However, in that experiment, salinities above 29 ppt failed to elicit an algal response. In another experiment, Symbiodiniaceae hosted by juvenile *Tridacna gigas* (giant clam) exhibited cell swelling, degradation, and pigment reductions at 18 ppt for 14 days, but algal cells within the clams were able to acclimatize to reduced salinity at 25 ppt (Maboloc et al. 2015). Thus, tank-based salinity stress experiments on *Orbicella* spp. holobionts from FGB can further confirm (or undermine) the conclusion that reduced salinity caused the gene expression changes we observed in *B. minutum* in the September 2017 samples. Regardless, our results inform our broader understanding of when and to what extent algal symbionts respond to changing environments and hosts.

### Transcriptomic Differences Between *Orbicella* Species

In the animal host, differences between congeneric coral species explained the most variation in expression (Figure 2–3A). Coral transcripts from both species represented in this study were mapped to the *O. faveolata* transcriptome and both species had similar mapping efficiencies to this reference. Previously classified as sister species within the genus *Montastraea*, *Orbicella faveolata* and *O. franksi* are largely sympatric (Weil and Knowlton 1994) and have many shared physical attributes that make them difficult to distinguish morphologically, though genetic variant analysis can resolve each species (Manzello et al. 2018). Differential gene expression between the two species that occurs independently of the effects of Hurricane Harvey are not the focus of this study, but do deserve consideration. The sequence datasets generated here can contribute to further research into species-specific coral expression, which, to our knowledge, has not been directly compared in these species. The top DEG between the two coral species shares substantial homology with an anemone (*Anthopleura asiatica*) toxin: DELTA-actitoxin-Aas1a (Kohno et al. 2009). This transcript, which was much more highly expressed in *O. faveolata* (LFC = 7.44, FDR = 3.98e-22), may indicate species-specific toxins that, to our knowledge, have yet to be characterized in these corals. We did not find any enriched GO categories between these coral species.

In *B. minutum*, coral host species had almost no effect on gene expression (Figure 2–3B). The one transcript that was differentially regulated between the two host species (LFC = 2.18, FDR = 1.03e-6) was unannotated in the transcriptome but shares sequence homology with an S-antigen protein (identity = 40.9%, E-value = 7.9e-68). Characterization of this protein is largely limited to variants associated with immune reactions in humans (*e.g*., Nussenblatt et al. 1982). Given the importance of host immune activation during the establishment and maintenance of symbiosis in corals (Mansfield et al. 2019) and the fact that these species have been found to host subtly different symbiont populations (Green et al. 2014), this differentially regulated transcript with antigenic potential deserves further investigation for its potential role in host–symbiont recognition.

#### Implications for Impacts of Future Storms on Reefs

The huge water surge that completely reduced salinity within Galveston Bay during Hurricane Harvey (Du et al. 2019) did not severely reduce salinity at the FGBNMS (Figure 1), probably because the water mass did not pass directly over the reef itself. Fortunately, the coral holobionts observed in this study were not exposed to extreme hyposalinity and the reduction in salinity that did occur was quickly alleviated. Reports of coral outcomes in past storms illustrate the potential damage hyposalinity can cause. In 1963, Hurricane Flora reduced coral reef salinity on several reefs in Eastern Jamaica to 3 ppt days after the storm and the region remained below 30 ppt for more than 5 weeks (Goreau 1964). As a result, multiple genera of corals in the region experienced substantial coral bleaching, though many colonies recovered fully within a few months. In 1987, heavy rains in Kaneohe Bay (Hawaii) substantially reduced salinity and caused mass coral mortality (Jokiel et al. 1993). Some species of corals (*e.g., Porites compressa*) in Kaneohe Bay recovered well, and comparisons to past mortality events support the ability of entire reefs to recover within 5–10 years if other stressors, such as pollution, are minimized. Coral reefs today suffer increasingly frequent stress events (Hughes et al. 2018). Our findings indicate that floodwaters following storms can trigger sub-lethal stress in corals; these impacts should be monitored and considered when assessing the cumulative threats to reef health.

## Conclusion

Though these corals survived the effects of Hurricane Harvey in the summer of 2017, a highly localized die-off event associated with hypoxic stress followed storm-driven flooding (2016 Tax Day Flood in Houston, TX, USA) on this reef the previous summer (Johnston et al. 2019; Kealoha *et al. in revision*). This event emphasizes the urgency to closely monitor the health of coral reefs subjected to multiple anthropogenic threats of increasing severity. Our genome-wide gene expression analysis of two coral species and their associated symbionts in the FGBNMS during the freshwater surge following Hurricane Harvey suggests that these endangered animals suffered sub-lethal stress, specifically related to redox state and mitochondrial function, which may compromise their ability to withstand subsequent stress (Adjeroud et al. 2009). Although these corals were able to recover, experimental evidence demonstrates that more extreme osmotic challenges can cause coral mortality and compromise photosynthetic function of their algal symbionts (Kerswell and Jones 2003; Downs et al. 2009). Monitoring coral health in the Gulf of Mexico is especially urgent considering the massive ongoing coral declines throughout the Caribbean (Rippe et al. 2019). Healthy coral colonies at the FGB sustain the local ecosystem and produce larvae that disperse throughout the Caribbean (Davies et al. 2017), which may help restore those devastated reefs.

## Conflict of Interest

The authors declare that the research was conducted in the absence of any commercial or financial relationships that could be construed as a potential conflict of interest.

## Author Contributions

RW analyzed gene expression and wrote the manuscript. SWD designed the experiment and contributed to the manuscript. AC contributed to the experimental design, collected the samples, and contributed to the manuscript. LQ prepared the gene expression libraries and contributed to the manuscript.

## Funding

This work was funded by NSF awards OCE-1800904 to SWD and OCE-1800914 to AC and stipend support for LQ was provided by a LINK Award through Brown University.

## Acknowledgments

September 2017 samples were collected on the R/V Manta and October 2017 samples were collected aboard the R/V Point Sur. We acknowledge the crew from these vessels and members of the NOAA FGBNMS office and Moody Gardens Aquarium. The authors particularly thank Marissa Nuttall, John Embesi and James MacMillan (NOAA), and Jake Emmert and Kaitlin Buhler (Moody Gardens) for field and logistical support, Emma Hickerson for facilitating collection permits, and Matthew Arnold for molecular work assistance.

## Notes

https://www.ncbi.nlm.nih.gov/bioproject/PRJNA552981

