## Supplemental Materials for "Gene expression of endangered coral (*Orbicella* spp.) in the Flower Garden Banks National Marine Sanctuary after Hurricane Harvey"

### *Supplementary Material*

**Supplemental Material Table 1**: Coral sampling information, including sampling time, location, and depth. "NR" indicates data not recorded.

| Sample Name | Time | Bank | Buoy | Lat Long | Species | Depth (m) |
|---|---|---|---|---|---|---|
| OFUE-1 | September 2017 | East | 1 | 27°52'54.84", 93°37'41.84" | *Orbicella faveolata* | NR |
| OFUE-2 | September 2017 | East | 1 | 27°52'54.84", 93°37'41.84" | *Orbicella faveolata* | NR |
| OFUE-3 | September 2017 | East | 1 | 27°52'54.84", 93°37'41.84" | *Orbicella faveolata* | NR |
| OFUE-4 | September 2017 | East | 1 | 27°52'54.84", 93°37'41.84" | *Orbicella faveolata* | NR |
| OFUE-5 | September 2017 | East | 1 | 27°52'54.84", 93°37'41.84" | *Orbicella faveolata* | NR |
| OFUE-6 | September 2017 | East | 1 | 27°52'54.84", 93°37'41.84" | *Orbicella faveolata* | NR |
| OFUE-7 | September 2017 | East | 1 | 27°52'54.84", 93°37'41.84" | *Orbicella faveolata* | NR |
| OFUE-8 | September 2017 | East | 1 | 27°52'54.84", 93°37'41.84" | *Orbicella faveolata* | NR |
| OFUE-9 | September 2017 | East | 1 | 27°52'54.84", 93°37'41.84" | *Orbicella faveolata* | NR |
| OFRUE-1 | September 2017 | East | 1 | 27°52'54.84", 93°37'41.84" | *Orbicella franksi* | NR |
| OFRUE-2 | September 2017 | East | 1 | 27°52'54.84", 93°37'41.84" | *Orbicella franksi* | NR |
| OFRUE-3 | September 2017 | East | 1 | 27°52'54.84", 93°37'41.84" | *Orbicella franksi* | NR |
| OFRUE-4 | September 2017 | East | 1 | 27°52'54.84", 93°37'41.84" | *Orbicella franksi* | NR |
| OFRUE-5 | September 2017 | East | 1 | 27°52'54.84", 93°37'41.84" | *Orbicella franksi* | NR |
| OFRUE-6 | September 2017 | East | 1 | 27°52'54.84", 93°37'41.84" | *Orbicella franksi* | NR |
| OFRUE-7 | September 2017 | East | 1 | 27°52'54.84", 93°37'41.84" | *Orbicella franksi* | NR |
| OFRUE-8 | September 2017 | East | 1 | 27°52'54.84", 93°37'41.84" | *Orbicella franksi* | NR |
| OFRUE-9 | September 2017 | East | 1 | 27°52'54.84", 93°37'41.84" | *Orbicella franksi* | NR |
| OFRUE-10 | September 2017 | East | 1 | 27°52'54.84", 93°37'41.84" | *Orbicella franksi* | NR |

| | | | | | | |
|---|---|---|---|---|---|---|
| OFUE-10 | September 2017 | East | 1 | 27°52'54.84", 93°37'41.84" | *Orbicella franksi* | NR |
| RR7-003 | October 2017 | East | 4 | 27°54'28.8", 93°36'0.72" | *Orbicella faveolata* | 20.4 |
| RR7-005 | October 2017 | East | 4 | 27°54'28.8", 93°36'0.72" | *Orbicella faveolata* | 21.3 |
| RR7-007 | October 2017 | East | 4 | 27°54'28.8", 93°36'0.72" | *Orbicella faveolata* | 21 |
| RR7-013 | October 2017 | East | 4 | 27°54'28.8", 93°36'0.72" | *Orbicella faveolata* | 19.2 |
| RR7-015 | October 2017 | East | 4 | 27°54'28.8", 93°36'0.72" | *Orbicella faveolata* | 19.2 |
| RR7-041 | October 2017 | East | 4 | 27°54'28.8", 93°36'0.72" | *Orbicella faveolata* | 20.4 |
| RR7-010 | October 2017 | West | 2 | 27°52'30.7194", 93°48'52.92" | *Orbicella faveolata* | 23.5 |
| RR7-019 | October 2017 | West | 2 | 27°52'30.7194", 93°48'52.92" | *Orbicella faveolata* | 22.6 |
| RR7-026 | October 2017 | West | 2 | 27°52'30.7194", 93°48'52.92" | *Orbicella faveolata* | 23.5 |
| RR7-028 | October 2017 | West | 2 | 27°52'30.7194", 93°48'52.92" | *Orbicella faveolata* | 22.6 |
| RR7-032 | October 2017 | West | 2 | 27°52'30.7194", 93°48'52.92" | *Orbicella faveolata* | 22.9 |
| RR7-033 | October 2017 | West | 2 | 27°52'30.7194", 93°48'52.92" | *Orbicella faveolata* | 24.1 |
| RR7-036 | October 2017 | West | 2 | 27°52'30.7194", 93°48'52.92" | *Orbicella faveolata* | 23.8 |
| RR7-004 | October 2017 | East | 4 | 27°54'28.8", 93°36'0.72" | *Orbicella franksi* | 21 |
| RR7-012 | October 2017 | East | 4 | 27°54'28.8", 93°36'0.72" | *Orbicella franksi* | 19.8 |
| RR7-016 | October 2017 | East | 4 | 27°54'28.8", 93°36'0.72" | *Orbicella franksi* | 19.2 |
| RR7-030 | October 2017 | East | 4 | 27°54'28.8", 93°36'0.72" | *Orbicella franksi* | 20.1 |
| RR7-037 | October 2017 | East | 4 | 27°54'28.8", 93°36'0.72" | *Orbicella franksi* | 19.8 |
| RR7-042 | October 2017 | East | 4 | 27°54'28.8", 93°36'0.72" | *Orbicella franksi* | 20.1 |
| RR7-009 | October 2017 | West | 2 | 27°52'30.7194", 93°48'52.92" | *Orbicella franksi* | 23.2 |
| RR7-017 | October 2017 | West | 2 | 27°52'30.7194", 93°48'52.92" | *Orbicella franksi* | 23.2 |
| RR7-020 | October 2017 | West | 2 | 27°52'30.7194", 93°48'52.92" | *Orbicella franksi* | 24.1 |

| | | | | | | |
|---|---|---|---|---|---|---|
| RR7-025 | October 2017 | West | 2 | 27°52'30.7194", 93°48'52.92" | *Orbicella franksi* | 23.8 |
| RR7-027 | October 2017 | West | 2 | 27°52'30.7194", 93°48'52.92" | *Orbicella franksi* | 23.8 |
| RR7-029 | October 2017 | West | 2 | 27°52'30.7194", 93°48'52.92" | *Orbicella franksi* | 23.5 |
| RR7-031 | October 2017 | West | 2 | 27°52'30.7194", 93°48'52.92" | *Orbicella franksi* | 23.2 |
| RR7-034 | October 2017 | West | 2 | 27°52'30.7194", 93°48'52.92" | *Orbicella franksi* | 23.8 |

**Supplemental Materials Table 2**: Coral host gene ontology enrichment. BP = biological processes; CC = cellular component; MF = molecular function. The direction indicates the direction of expression change of the genes in that category. Samples were collected either during a period of 'sub-lethal stress' (September 2017) or 'recovery' (October 2017).

| Comparison | GO Category | Direction | P.adjusted | Name | Term |
|---|---|---|---|---|---|
| Collection Time | BP | Up in Recovery | 3.42E-03 | organelle fission | GO:0000280;GO:0048285 |
| Collection Time | BP | Up in Recovery | 2.27E-02 | RNA processing | GO:0006396 |
| Collection Time | BP | Up in Recovery | 3.42E-03 | mitosis | GO:0007067 |
| Collection Time | BP | Up in Recovery | 2.77E-02 | chromatin organization | GO:0016569;GO:0016568; GO:0006325 |
| Collection Time | BP | Up in Recovery | 1.05E-04 | cell cycle process | GO:0022402 |
| Collection Time | BP | Up in Recovery | 4.89E-02 | regulation of organelle organization | GO:0033043 |
| Collection Time | BP | Up in Recovery | 2.55E-02 | Wnt signaling pathway involved in dorsal/ventral axis specification | GO:0044332 |
| Collection Time | BP | Up in Recovery | 1.34E-02 | regulation of RNA export from nucleus | GO:0046831;GO:0032239 |
| Collection Time | BP | Up in Recovery | 1.01E-03 | chromosome organization | GO:0051276 |
| Collection Time | BP | Up in Recovery | 3.42E-03 | cell division | GO:0051301 |
| Collection Time | BP | Up in Recovery | 4.83E-02 | positive regulation of protein transport | GO:0090316;GO:0032388; GO:0051222 |
| Collection Time | BP | Up in Sub-Lethal Stress | 4.09E-03 | translation | GO:0006412 |
| Collection Time | BP | Up in Sub-Lethal Stress | 4.89E-02 | cognition | GO:0007611;GO:0050890 |
| Collection Time | BP | Up in Sub-Lethal Stress | 2.27E-02 | nucleoside triphosphate biosynthetic process | GO:0009201;GO:0009142; GO:0009206;GO:0009145 |

| | | | | | |
|---|---|---|---|---|---|
| Collection Time | BP | Up in Sub-Lethal Stress | 3.84E-02 | extracellular matrix organization | GO:0030198;GO:0043062 |
| Collection Time | BP | Up in Sub-Lethal Stress | 2.55E-02 | multicellular organismal metabolic process | GO:0032963;GO:0044259; GO:0044236;GO:0030574; GO:0044243 |
| Collection Time | BP | Up in Sub-Lethal Stress | 3.86E-02 | cell redox homeostasis | GO:0045454 |
| Collection Time | CC | Up in Recovery | 4.15E-02 | exocyst | GO:0000145 |
| Collection Time | CC | Up in Recovery | 4.73E-02 | ubiquitin ligase complex | GO:0000151 |
| Collection Time | CC | Up in Recovery | 4.20E-02 | nuclear chromosome | GO:0000228 |
| Collection Time | CC | Up in Recovery | 4.83E-02 | chromosome, centromeric region | GO:0000775 |
| Collection Time | CC | Up in Recovery | 2.13E-02 | chromatin | GO:0000785 |
| Collection Time | CC | Up in Recovery | 1.32E-02 | spindle pole | GO:0000922 |
| Collection Time | CC | Up in Recovery | 4.15E-02 | chromosome | GO:0005694 |
| Collection Time | CC | Up in Recovery | 1.27E-02 | centrosome | GO:0005813 |
| Collection Time | CC | Up in Recovery | 1.30E-03 | microtubule organizing center | GO:0005815 |
| Collection Time | CC | Up in Recovery | 3.87E-02 | spindle | GO:0005819 |
| Collection Time | CC | Up in Recovery | 4.15E-02 | myofilament | GO:0005865;GO:0036379 |
| Collection Time | CC | Up in Recovery | 3.10E-03 | microtubule | GO:0005874 |
| Collection Time | CC | Up in Recovery | 1.41E-03 | nuclear body | GO:0016604 |
| Collection Time | CC | Up in Recovery | 9.04E-03 | nuclear speck | GO:0016607 |
| Collection Time | CC | Up in Recovery | 3.66E-02 | cell body | GO:0043025;GO:0044297 |
| Collection Time | CC | Up in Recovery | 5.58E-04 | chromosomal part | GO:0044427 |
| Collection Time | CC | Up in Recovery | 7.09E-06 | cytoskeletal part | GO:0044430 |
| Collection Time | CC | Up in Recovery | 3.53E-04 | nucleoplasm part | GO:0044451 |

| | | | | | |
|---|---|---|---|---|---|
| Collection Time | CC | Up in Recovery | 4.54E-02 | intermediate filament cytoskeleton | GO:0045111 |
| Collection Time | CC | Up in Recovery | 1.30E-03 | neuron part | GO:0097458 |
| Collection Time | CC | Up in Sub-Lethal Stress | 1.10E-04 | extracellular region | GO:0005576 |
| Collection Time | CC | Up in Sub-Lethal Stress | 4.20E-02 | collagen trimer | GO:0005581 |
| Collection Time | CC | Up in Sub-Lethal Stress | 4.20E-02 | extracellular space | GO:0005615 |
| Collection Time | CC | Up in Sub-Lethal Stress | 2.36E-03 | respiratory chain complex I | GO:0005747;GO:0045271;GO:0030964 |
| Collection Time | CC | Up in Sub-Lethal Stress | 2.73E-02 | endoplasmic reticulum lumen | GO:0005788 |
| Collection Time | CC | Up in Sub-Lethal Stress | 1.10E-04 | endoplasmic reticulum membrane | GO:0005789 |
| Collection Time | CC | Up in Sub-Lethal Stress | 9.83E-03 | ribosome | GO:0005840 |
| Collection Time | CC | Up in Sub-Lethal Stress | 4.20E-02 | proton-transporting two-sector ATPase complex | GO:0016469 |
| Collection Time | CC | Up in Sub-Lethal Stress | 5.58E-04 | extracellular matrix | GO:0031012;GO:0005578 |
| Collection Time | CC | Up in Sub-Lethal Stress | 3.60E-02 | integral component of mitochondrial outer membrane | GO:0031307 |
| Collection Time | CC | Up in Sub-Lethal Stress | 1.27E-02 | myelin sheath | GO:0043209 |
| Collection Time | CC | Up in Sub-Lethal Stress | 4.94E-02 | ribosomal subunit | GO:0044391 |
| Collection Time | CC | Up in Sub-Lethal Stress | 6.89E-08 | endoplasmic reticulum part | GO:0044432 |
| Collection Time | CC | Up in Sub-Lethal Stress | 4.04E-07 | mitochondrial membrane part | GO:0044455 |
| Collection Time | CC | Up in Sub-Lethal Stress | 3.59E-02 | membrane region | GO:0098589 |
| Collection Time | MF | Up in Recovery | 3.82E-02 | chromatin binding | GO:0003682 |

| | | | | | |
|---|---|---|---|---|---|
| Collection Time | MF | Up in Recovery | 1.77E-02 | motor activity | GO:0003774 |
| Collection Time | MF | Up in Recovery | 1.08E-02 | actin binding | GO:0003779 |
| Collection Time | MF | Up in Recovery | 2.11E-02 | helicase activity | GO:0004386 |
| Collection Time | MF | Up in Recovery | 5.97E-03 | ligase activity, forming carbon-nitrogen bonds | GO:0004842;GO:0019787;GO:0016881;GO:0016879 |
| Collection Time | MF | Up in Recovery | 3.36E-02 | guanyl-nucleotide exchange factor activity | GO:0005085 |
| Collection Time | MF | Up in Recovery | 5.82E-03 | nucleoside-triphosphatase regulator activity | GO:0005096;GO:0030695;GO:0060589 |
| Collection Time | MF | Up in Recovery | 3.69E-03 | ligase activity | GO:0016874 |
| Collection Time | MF | Up in Recovery | 5.82E-03 | ATPase activity | GO:0016887 |
| Collection Time | MF | Up in Recovery | 3.49E-02 | GTPase binding | GO:0017016;GO:0031267;GO:0051020 |
| Collection Time | MF | Up in Recovery | 5.82E-03 | hydrolase activity, acting on acid anhydrides | GO:0017111;GO:0016462;GO:0016818;GO:0016817 |
| Collection Time | MF | Up in Recovery | 2.35E-02 | Wnt-protein binding | GO:0017147 |
| Collection Time | MF | Up in Recovery | 4.31E-02 | enzyme binding | GO:0019899 |
| Collection Time | MF | Up in Recovery | 4.05E-02 | heat shock protein binding | GO:0031072 |
| Collection Time | MF | Up in Recovery | 2.02E-02 | Wnt-activated receptor activity | GO:0042813 |
| Collection Time | MF | Up in Recovery | 2.75E-02 | mitogen-activated protein kinase binding | GO:0051019 |
| Collection Time | MF | Up in Sub-Lethal Stress | 3.69E-03 | structural constituent of ribosome | GO:0003735 |
| Collection Time | MF | Up in Sub-Lethal Stress | 2.32E-02 | endopeptidase activity | GO:0004175 |

| | | | | | |
|---|---|---|---|---|---|
| Collection Time | MF | Up in Sub-Lethal Stress | 3.69E-03 | metalloendopeptidase activity | GO:0004222 |
| Collection Time | MF | Up in Sub-Lethal Stress | 2.75E-02 | oxidoreductase activity, acting on peroxide as acceptor | GO:0004601;GO:0016684 |
| Collection Time | MF | Up in Sub-Lethal Stress | 4.31E-02 | structural molecule activity | GO:0005198 |
| Collection Time | MF | Up in Sub-Lethal Stress | 1.72E-02 | metallopeptidase activity | GO:0008237 |
| Collection Time | MF | Up in Sub-Lethal Stress | 1.08E-02 | monovalent inorganic cation transmembrane transporter activity | GO:0015077 |
| Collection Time | MF | Up in Sub-Lethal Stress | 2.11E-02 | hydrogen ion transmembrane transporter activity | GO:0015078 |
| Collection Time | MF | Up in Sub-Lethal Stress | 1.35E-02 | antioxidant activity | GO:0016209 |
| Collection Time | MF | Up in Sub-Lethal Stress | 5.82E-03 | oxidoreductase activity | GO:0016491 |

**Supplemental Materials Table 3**: Algal symbiont *Breviolum minutum* gene ontology enrichment. BP = biological processes; CC = cellular component; MF = molecular function; *O. faveolata* = *Orbicella faveolata*. The direction indicates the direction of expression change of the genes in that category. Samples were collected either during a period of 'sub-lethal stress' (September 2017) or 'recovery' (October 2017).

| Comparison | GO Category | Direction | P.adjusted | Name | Term |
|---|---|---|---|---|---|
| Collection Time | BP | Up in Recovery | 2.20E-03 | RNA splicing, via transesterification reactions | GO:0000398;GO:0000377;GO:0000375 |
| Collection Time | BP | Up in Recovery | 1.58E-02 | rRNA metabolic process | GO:0006364;GO:0016072 |
| Collection Time | BP | Up in Recovery | 7.84E-10 | RNA processing | GO:0006396 |
| Collection Time | BP | Up in Recovery | 6.48E-08 | mRNA metabolic process | GO:0006397;GO:0016071 |
| Collection Time | BP | Up in Recovery | 6.08E-07 | RNA splicing | GO:0008380 |
| Collection Time | BP | Up in Recovery | 8.18E-04 | ncRNA metabolic process | GO:0034470;GO:0034660 |
| Collection Time | BP | Up in Recovery | 1.06E-02 | ribonucleoprotein complex biogenesis | GO:0042254;GO:0022613 |
| Collection Time | BP | Up in Recovery | 8.87E-03 | cellular component biogenesis | GO:0044085 |
| Collection Time | BP | Up in Sub-Lethal Stress | 2.32E-02 | steroid biosynthetic process | GO:0006694 |
| Collection Time | BP | Up in Sub-Lethal Stress | 1.58E-02 | ion transport | GO:0006811 |
| Collection Time | BP | Up in Sub-Lethal Stress | 4.95E-02 | single-organism catabolic process | GO:0044712 |
| Collection Time | BP | Up in Sub-Lethal Stress | 2.76E-05 | transmembrane transport | GO:0055085 |
| Collection Time | CC | Up in Recovery | 3.99E-03 | spliceosomal complex | GO:0005681 |
| Collection Time | CC | Up in Recovery | 2.71E-02 | nucleolus | GO:0005730 |
| Collection Time | CC | Up in Recovery | 1.51E-04 | ribonucleoprotein complex | GO:0030529 |
| Collection Time | CC | Up in Recovery | 2.71E-02 | catalytic step 2 spliceosome | GO:0071013 |
| Collection Time | CC | Up in Sub-Lethal Stress | 2.71E-02 | oxidoreductase complex | GO:1990204 |
| Collection Time | MF | Up in Recovery | 1.82E-03 | DNA binding | GO:0003677 |
| Collection Time | MF | Up in Recovery | 1.85E-09 | RNA binding | GO:0003723 |

| | | | | | |
|---|---|---|---|---|---|
| Collection Time | MF | Up in Recovery | 7.05E-04 | helicase activity | GO:0004386 |
| Collection Time | MF | Up in Recovery | 4.34E-04 | ATP-dependent helicase activity | GO:0008026;GO:0070035 |
| Collection Time | MF | Up in Recovery | 2.87E-02 | DNA-dependent ATPase activity | GO:0008094 |
| Collection Time | MF | Up in Recovery | 2.87E-02 | transferase activity, transferring phosphorus-containing groups | GO:0016772 |
| Collection Time | MF | Up in Recovery | 4.55E-02 | ATPase activity | GO:0016887;GO:0042623 |
| Collection Time | MF | Up in Recovery | 3.16E-03 | pyrophosphatase activity | GO:0017111;GO:0016462;GO:0016818;GO:0016817 |
| Collection Time | MF | Up in Sub-Lethal Stress | 4.95E-02 | monooxygenase activity | GO:0004497 |
| Collection Time | MF | Up in Sub-Lethal Stress | 2.06E-03 | transporter activity | GO:0005215 |
| Collection Time | MF | Up in Sub-Lethal Stress | 3.21E-02 | oxidoreductase activity | GO:0016709 |
| Collection Time | MF | Up in Sub-Lethal Stress | 3.93E-03 | transmembrane transporter activity | GO:0022857;GO:0015075;GO:0022891;GO:0022892 |
| Host Species | BP | Up in *O. faveolata* | 3.58E-02 | RNA splicing, via transesterification reactions | GO:0000398;GO:0000377;GO:0000375 |
| Host Species | BP | Up in *O. faveolata* | 3.58E-02 | RNA processing | GO:0006396 |
| Host Species | BP | Up in *O. faveolata* | 1.66E-02 | mRNA metabolic process | GO:0006397;GO:0016071 |
| Host Species | BP | Up in *O. faveolata* | 2.55E-04 | RNA splicing | GO:0008380 |
| Host Species | BP | Up in *O. faveolata* | 1.66E-02 | cell cycle process | GO:0022402 |
| Host Species | CC | Up in *O. faveolata* | 1.69E-02 | spliceosomal complex | GO:0005681 |
| Host Species | CC | Up in *O. faveolata* | 3.80E-02 | ribonucleoprotein complex | GO:0030529 |
